# DelPi Learns Generalizable Peptide–Signal Correspondence for Mass Spectrometry-Based Proteomics

**DOI:** 10.64898/2026.01.06.697814

**Authors:** Jungkap Park, Kyunggon Kim, Un-Beom Kang, Sangtae Kim

## Abstract

Peptide identification in mass spectrometry–based proteomics has traditionally relied on handcrafted features or simplified probabilistic approaches that limit the interpretation of structured peptide evidence. We present DelPi, an open-source peptide identification framework that learns generalizable peptide–signal correspondence from raw spectra through self-supervised pre-training followed by task-specific fine-tuning. With model distillation enabling practical deployment, DelPi expands the interpretation of peptide evidence across data-independent and data-dependent acquisition while maintaining robust false discovery control.

---

Deep representation learning has transformed data interpretation by replacing feature engineering with end-to-end models trained on raw data^1,2^. In mass spectrometry–based proteomics, however, widely used identification tools for both data-dependent acquisition (DDA) and data-independent acquisition (DIA) remain dependent on handcrafted features or simplified scoring schemes^3–8^. Recent deep learning approaches introduce data-driven elements, but retain feature-based scoring^6^ or rely on computationally intensive, library-dependent designs^9^, thereby failing to substantially expand the interpretation of complex peptide evidence without sacrificing practical deployment. For example, DIA-BERT introduces a feature-free Transformer-based scorer^9^ but is restricted to DIA and library-based workflows, relies on supervised pre-training that is not designed to capture intrinsic peptide-derived signal structure, and requires expensive per-run fine-tuning, limiting both generalizability and scalability. In addition, the proprietary or closed-source nature of several major tools such as DIA-NN^3^ and MSFragger^7^ limits transparency and rapid adaptation to new experimental settings.

Here we present DelPi, an open-source peptide identification framework built on a pre-trained Transformer encoder that combines improved interpretation of complex peptide evidence with practical deployment on a consumer-grade GPU (Fig. 1). Using self-supervised learning, DelPi learns a generalizable representation of peptide-derived signals directly from raw spectra, removing reliance on handcrafted features and supporting unified peptide identification across both DIA and DDA. The trained encoder is further distilled into a compact student model to enable efficient inference.

**Figure 1.**
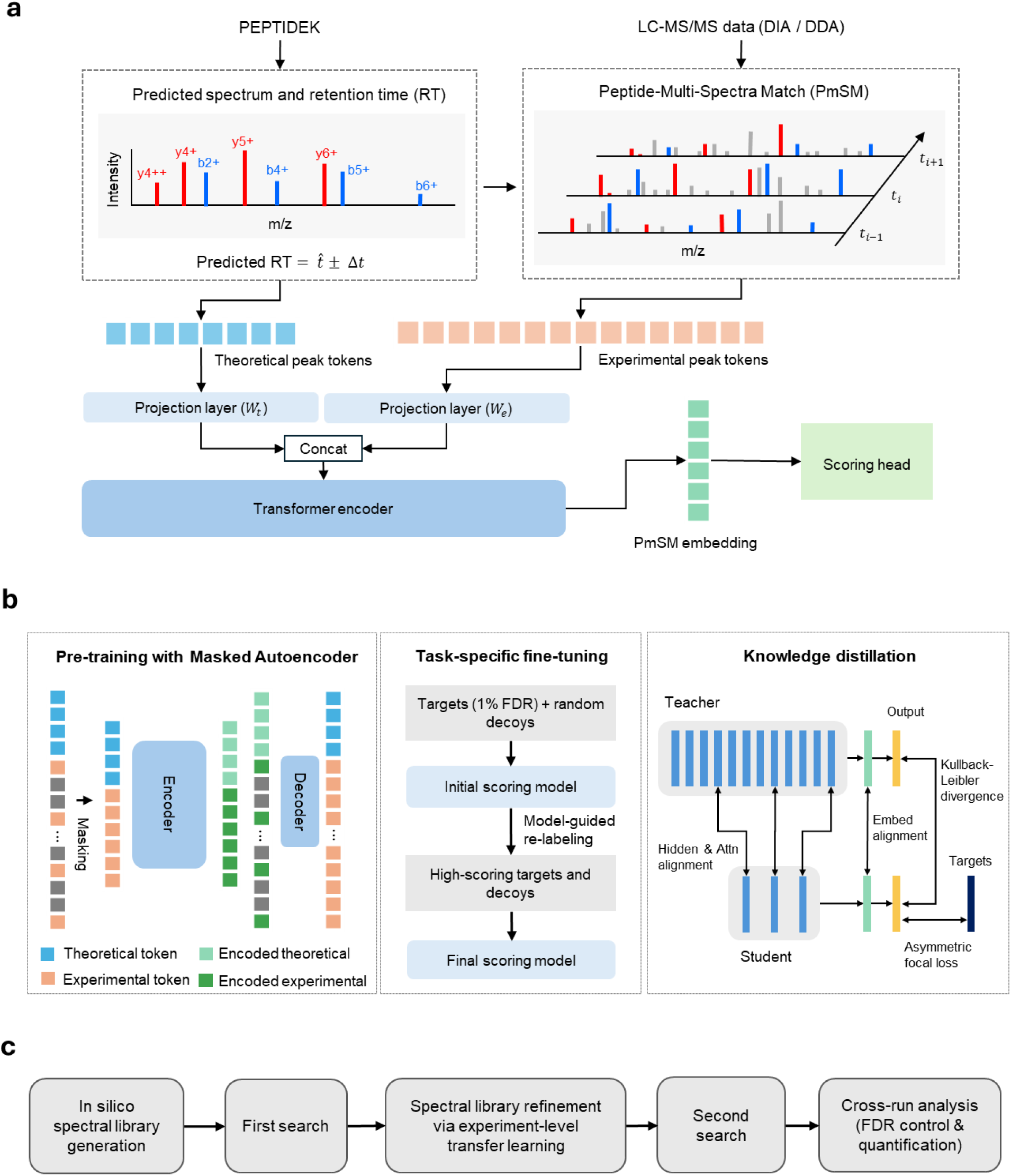
Overview of the DelPi framework. **a,** Model architecture. Each candidate precursor is represented as a Peptide–Multi–Spectra Match (PmSM), which aggregates theoretical precursor and fragment-ion peaks with experimentally observed peaks across the peptide elution profile, preserving precursor isotope envelopes, fragment-ion ladders, chromatographic co-elution, and elution-dependent intensity changes. Theoretical and experimental peaks are tokenized and processed by a Transformer encoder to produce a latent embedding, which is mapped to a likelihood score by a two-layer fully connected scoring head. **b,** Training pipeline. The training strategy comprises three stages. In self-supervised masked autoencoder (MAE) pre-training, the encoder learns a generalizable representation of peptide-derived signals by reconstructing masked experimental peak tokens. Task-specific fine-tuning then optimizes the pre-trained encoder for peptide identification using target and decoy PmSMs. The training set is progressively expanded by incorporating high-confidence predictions, allowing the model to learn discriminative patterns beyond those captured by conventional tools. Finally, the 12-layer teacher model is distilled into a compact 3-layer student model by aligning predictions and internal representations for efficient inference. **c,** Search workflow. DelPi performs an initial search using an in silico spectral library to identify high-confidence PmSMs, which are used for experiment-level transfer learning to adapt fragment-intensity and retention-time predictors. The adapted predictors generate a refined library for a second search, followed by cross-run scoring, quantification, and protein inference.

Central to DelPi is the Peptide–Multi–Spectra Match (PmSM), a unified representation that aggregates theoretical peptide features with experimentally observed signals across the elution profile (Fig. 1a). Unlike traditional peptide–spectrum matches (PSMs), which associate a peptide with a single spectrum, a PmSM represents a peptide elution event as a single learning unit. Each PmSM is tokenized into a sequence of theoretical and experimental peak tokens, enabling a Transformer model to align theoretical ions with observed peaks and to capture coherent chromatographic and spectral structure.

To learn a generalizable representation of PmSMs, DelPi adopts the Masked Autoencoder (MAE) pre-training strategy (Fig. 1b)^10^. Signals arising from the same peptide exhibit structured redundancy across scans, isotopes, and fragment ions. During pre-training, a large fraction of experimental peak tokens is masked and reconstructed from theoretical and remaining experimental tokens, discouraging trivial interpolation and encouraging the encoder to learn structured dependencies. Although pre-training was performed exclusively on DIA datasets, the learned representations generalized effectively to DDA, indicating that the encoder learns intrinsic patterns of peptide-derived signals across instruments and acquisition modes.

The pre-trained encoder is then specialized for peptide identification using an iterative semi-supervised fine-tuning scheme (Fig. 1b). Fine-tuning is initialized from PmSMs corresponding to 1% false discovery rate (FDR) targets identified by conventional search tools, together with decoy-derived PmSMs, and progressively expanded by incorporating high-confidence predictions. This design enables the model to refine peptide discrimination beyond the fixed, manually defined feature space of conventional approaches. An asymmetric focal loss^11^ down-weights noisy positives while emphasizing informative hard negatives, improving discrimination of peptide evidence at stringent FDR thresholds. The fine-tuned 12-layer encoder is distilled into a compact 3-layer student model, substantially reducing inference time with minimal loss of accuracy (Fig. 1b).

DelPi integrates the learned representation into a two-stage search workflow analogous to modern DIA^3,6^ and iterative DDA^12^ strategies (Fig. 1c). An in silico spectral library is generated using deep learning–based predictors of fragment-ion intensities and peptide retention times. In the first stage, high-confidence peptides are identified and used for experiment-level transfer learning to adapt the models for spectral library generation to instrument and chromatographic conditions. The adapted models are then used to construct a refined spectral library for a second, more sensitive search. Results are consolidated across runs through global scoring, retention-time alignment, and protein inference.

We evaluated DelPi on six public datasets covering diverse instruments, acquisition modes, and sample types: four DIA datasets (mixed-species dilution-series^13^, single-cell proteomics^14^, neat plasma^15^, and phosphoproteomics^16^) and two DDA datasets (global proteomics^17^ and TMT-labeled phosphoproteomics^18^) from CPTAC studies (Supplementary Table 1). For DIA benchmarks, DelPi was compared with DIA-NN^3^, AlphaDIA^6^, and DIA-BERT^9^, whereas for DDA benchmarks it was compared with MS-GF+^4^ and Sage^5^. An earlier version of DIA-NN (v1.8.1) was used and MSFragger^7^ was not included because newer versions of DIA-NN and MSFragger require separate licenses for use by commercial organizations. FDR control was assessed using entrapment-based false discovery proportion (FDP) estimates, reporting a lower-bound FDP, defined as the fraction of entrapment identifications among reported targets, and a combined FDP adjusted by the entrapment-to-target database ratio^19^.

Across DIA datasets, DelPi achieved improved identification sensitivity with well-controlled false discovery behavior (Fig. 2a and Supplementary Fig. 2). In the mixed-species dilution-series benchmark, DelPi identified 24-30% more precursors and 11-22% more protein groups than DIA-NN in both single-run and multi-run analyses, while exhibiting comparable combined FDPs. In the neat plasma and single-cell datasets, DIA-NN occasionally reported slightly more identifications, but these gains were accompanied by higher FDPs, whereas DelPi maintained lower FDPs. A similar pattern was observed in the phosphoproteomics dataset, where DelPi identified more than twice as many precursors and 26% more protein groups than DIA-NN, while maintaining comparable or lower FDPs. In contrast, AlphaDIA often reported higher identification counts but exhibited substantially elevated FDPs.

**Figure 2.**
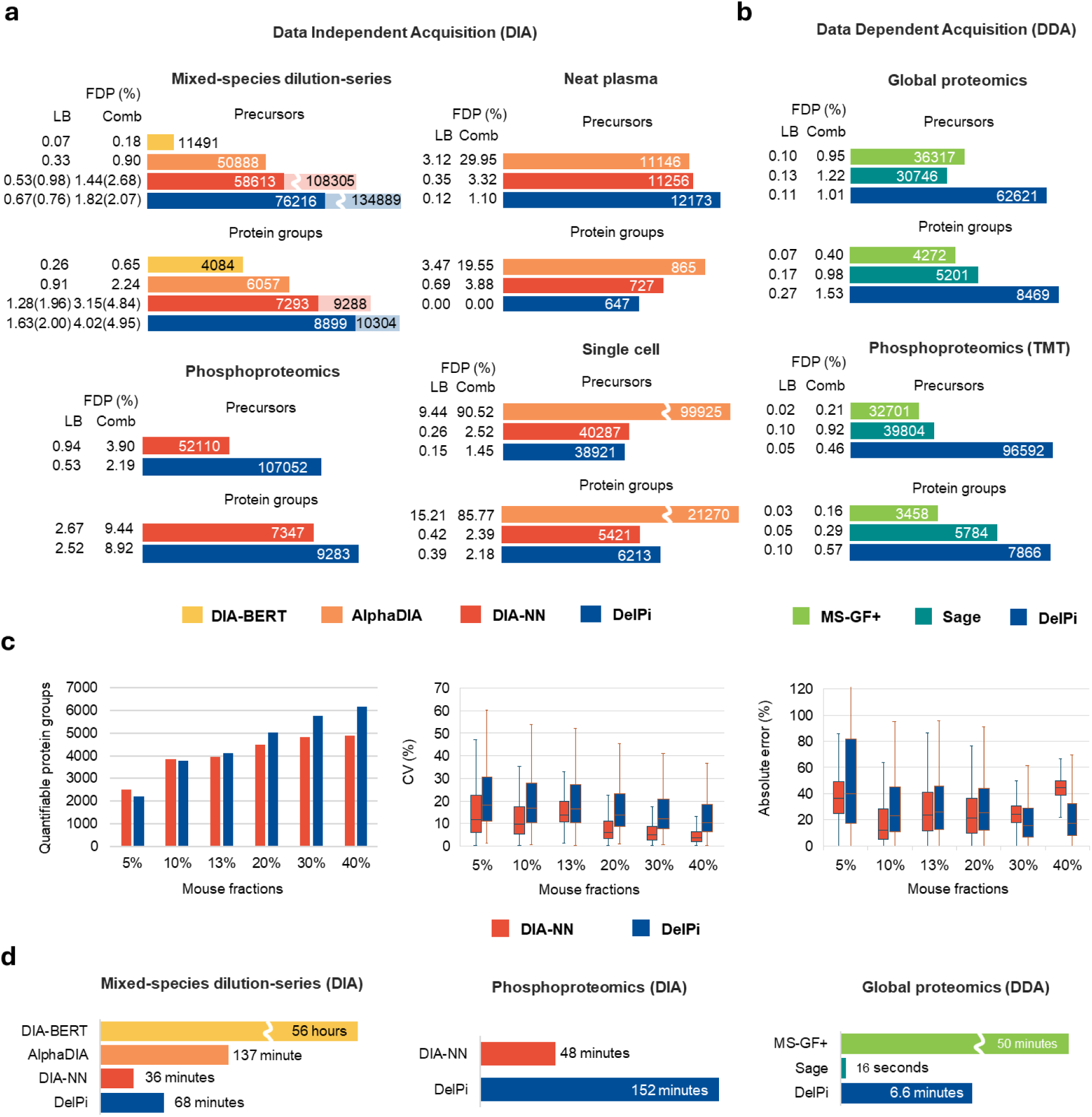
Benchmark results. **a,** Identification performance on DIA datasets. Bar charts show the numbers of identified precursors and protein groups at 1% false discovery rate (FDR), together with the entrapment-based lower-bound (LB) false discovery proportion (FDP) and the combined FDP (Comb). For the mixed-species dilution-series dataset, saturated bars indicate single-run analysis, whereas light bars indicate multi-run analysis (all 35 runs for DIA-NN and DelPi); corresponding FDPs are shown in parentheses. **b,** Identification performance on DDA datasets. **c,** Quantification performance on the mixed-species dilution-series DIA dataset, consisting of mouse-yeast mixtures with defined mouse fractions (5–40%). Plots show the numbers of quantified protein groups, boxplots of coefficients of variation (CVs) and relative absolute errors across mixing ratios. **d,** Runtime comparison. Average processing time per run across representative DIA and DDA benchmark datasets.

In the DDA benchmarks, DelPi substantially outperformed traditional PSM-centric search engines while maintaining comparable false discovery control (Fig. 2b and Supplementary Fig. 3). Compared with MS-GF+, DelPi identified 72–195% more precursors and 98–127% more protein groups, and compared with Sage it identified 103–142% more precursors and 36–63% more protein groups. Although Sage supports chimeric searches for identifying multiple peptides per spectrum (enabled here) and showed modest gains over MS-GF+, DelPi exhibited substantially higher sensitivity than both tools, highlighting the advantage of PmSM-based representation learning over conventional PSM-centric, feature-based scoring.

Quantification on the mixed-species dilution-series DIA dataset showed that DelPi quantified a similar or greater number of spiked-in mouse protein groups than DIA-NN across all mixing ratios (Fig. 2c). DelPi exhibited higher variability and larger absolute ratio errors, particularly at low mixing ratios, reflecting the simpler nature of its current label-free quantification approach compared with the more advanced interference-aware algorithms implemented in DIA-NN. Further improvements in DelPi’s label-free quantification are left for future work.

Runtime benchmarks showed that while DelPi was slower than traditional tools such as DIA-NN or Sage (Fig. 2d), it was substantially faster than other deep learning–based approaches, including AlphaDIA and DIA-BERT. For a typical 25-minute Orbitrap Astral liquid chromatography–mass spectrometry (LC–MS) analysis from a human sample, DelPi completed analysis in approximately 20 minutes on a single consumer-grade GPU (for example, an RTX 4090), supporting practical laboratory use.

Together, these results establish representation learning as a practical foundation for peptide identification applicable to both DIA and DDA workflows with reliable false discovery control. DelPi provides a unified, acquisition-agnostic identification framework that enables routine use in standard laboratory environments. Its open and extensible implementation enables future extensions to downstream analyses, such as interference-aware DIA quantification, post-translational modification localization, and robust cross-run integration, establishing a clear pathway toward scalable proteomics studies across diverse experimental settings.

## Online Methods

### Model architecture

#### Peptide–Multi–Spectra Match (PmSM)

For each candidate precursor, defined as a modified peptide sequence with a specific charge state, DelPi constructs a Peptide–Multi–Spectra Match (PmSM), a structured data object that aggregates MS1 and MS2 evidence across a precursor elution event.

Theoretical peaks corresponding to a candidate precursor are matched to observed peaks in MS1 and MS2 spectra using a mass tolerance of 10 ppm, applied consistently during training, search, and benchmarking. The PmSM is constructed from the theoretical peaks and the matched experimental peaks. For the precursor ion, up to three isotopic peaks are matched when present, and for each theoretical fragment ion, up to two corresponding observed isotopic peaks are matched. Theoretical peaks are obtained from in silico spectral library prediction as described below.

#### Tokenization

Each PmSM is tokenized into a sequence of numerical tokens encoding both theoretical and experimental peaks. The concatenated sequence of theoretical and experimental peak tokens serves as the input to the Transformer encoder for precursor-level scoring.

##### Theoretical peak tokens

Theoretical peak tokens encode expected precursor isotopes and fragment-ion peaks. Each token encodes predicted m/z, predicted intensity, charge state, isotope index, ion type (precursor, b ion, or y ion), and fragment index (for example, 5 for a y5 ion). DelPi includes the three most intense precursor isotopes and the sixteen most intense predicted fragment ions, yielding a total of 19 theoretical peak tokens per PmSM.

##### Experimental peak tokens

Experimental peak tokens encode observed signals and associated metadata. Each token encodes mass error, normalized intensity, retention-time index, parametric intensity score, and metadata including charge state, isotope index, ion type, and fragment index. Mass error is computed in ppm with sign defined as (observed m/z − theoretical m/z) and is then divided by the 10, yielding a rescaled value in [−1, 1]. Intensities are normalized separately for precursor and fragment peaks such that the maximum intensity within each group is 1.0. The retention-time index is a zero-based integer reflecting the temporal order of scans within each PmSM. The parametric intensity score represents the relative position of each peak intensity within its spectrum and is computed by fitting a Gaussian distribution to the spectrum’s raw peak intensities and encoding the cumulative distribution function (CDF) value (range 0–1).

#### Transformer-based PmSM encoder and scoring head

All tokens are linearly projected into a shared embedding space. Token-specific attributes encoding temporal or sequential order, including the retention-time index and fragment index, are encoded using sinusoidal positional encodings and added to the token embeddings. The resulting token sequence is processed by a 12-layer Transformer encoder with an embedding dimension of 192 and 12 attention heads. Multi-head self-attention integrates predicted and observed fragmentation patterns and multi-scan chromatographic information into a latent PmSM embedding that encodes correspondence to theoretical expectations as well as observed signal structure.

A lightweight multilayer perceptron (MLP) scoring head is attached to the PmSM encoder to perform binary classification of true and false PmSMs. The scoring head consists of two fully connected layers with 64 and 32 neurons, respectively, and maps the encoder embedding to a scalar logit. This logit is transformed using a sigmoid function to yield a likelihood score between 0 and 1, representing the model’s confidence that a given PmSM corresponds to a true match.

### Training procedures

MAE-based pre-training

#### Training data

High-confidence target identifications were collected from DIA training datasets (Supplementary Table 2) using AlphaDIA, DIA-NN, EncyclopeDIA, and Sage. For each reported target precursor at 1% FDR, a corresponding PmSM was constructed using the same PmSM extraction procedures described in Search workflow. In total, approximately 7.9 million target PmSMs were collected for pre-training.

#### Objective

DelPi adapted a masked autoencoder (MAE) objective to tokenized PmSMs. A masking ratio of 0.75 was applied to experimental peak tokens. The model was trained to reconstruct masked experimental-token fields from visible experimental tokens together with theoretical tokens. Reconstruction targets included rescaled mass error, normalized intensity, and peak intensity percentile, optimized with an unweighted mean-squared-error (MSE) loss over these three fields. The MAE decoder had 8 layers with a 128-dimensional embedding, whereas the encoder had 12 layers with a 192-dimensional embedding.

#### Optimization

Pre-training was performed for 210 epochs with a batch size of 512 using the AdamW optimizer, a base learning rate of 1 × 10⁻³, and a weight decay of 0.05. A cosine learning-rate schedule with warm-up and restarts was used, with warm-up for the first 10 epochs and learning-rate restarts every 40 epochs.

##### Task-specific fine-tuning

#### Training data

After MAE pre-training, the encoder was fine-tuned with an attached MLP scoring head to classify target and decoy PmSMs. Target PmSMs were derived from 1% FDR target identifications reported by established search engines (DIA: AlphaDIA, DIA-NN, EncyclopeDIA, and Sage; DDA: MS-GF+ and Sage). Decoy PmSMs were generated using a pseudo-reverse strategy (reversing peptide sequences while preserving the C-terminal residue and amino acid composition). PmSMs were retained only if they met minimum matched-peak criteria (≥12 matched peaks for DIA; ≥3 for DDA). The initial training set comprised ∼7.9 million DIA targets and 427 million DIA decoys, and ∼4.8 million DDA targets and 104 million DDA decoys.

#### Objective

Fine-tuning used an asymmetric focal loss^11^. To reduce sensitivity to potential label noise among positives, the positive focusing parameter was set to 0. To prevent optimization from being dominated by abundant easy negatives and to emphasize informative hard negatives, the negative focusing parameter was set to 4.

#### Two-stage refinement

Fine-tuning was performed in two stages. In Stage 1, the encoder was fine-tuned on the initial labeled set. The resulting model was used to rescore candidate PmSMs extracted from raw mass spectrometry data. High-confidence predictions were added to the training pool (likelihood >0.9 for targets and >0.5 for decoys), conservatively expanding positives while broadening negative coverage. This expansion added ∼14 million DIA targets and 51 million DIA decoys, and ∼5.5 million DDA targets and 4.2 million DDA decoys. In Stage 2, the encoder was fine-tuned on the expanded dataset. Separate fine-tuned models were trained for DIA and DDA.

#### Optimization

Fine-tuning was performed for 40 epochs per stage using AdamW with a base learning rate of 1×10⁻⁴ and weight decay of 0.01. In each epoch, 40 million examples were randomly down-sampled. The positive-to-negative sampling ratio was 1:5 in Stage 1 and 1:3 in Stage 2. In Stage 2 down-sampling, the ratio of high-scoring decoy PmSMs to randomly sampled decoy PmSMs was kept at 1:2 for DIA and 1:6 for DDA.

##### Knowledge Distillation

The 12-layer fine-tuned teacher model was distilled into a compact 3-layer student model using the expanded training datasets from Stage 2 fine-tuning. Distillation employed a multi-objective loss combining (i) the asymmetric focal task loss against ground-truth labels, (ii) Kullback–Leibler divergence between softened logits (temperature τ = 2.0), and (iii) MSE losses aligning the final embeddings, intermediate hidden states, and attention maps of the teacher and student models. The task loss and embedding-alignment loss were each weighted at 1.0, the KL-divergence term at 0.3, the hidden-state alignment loss at 0.5, and the attention-map alignment loss at 0.25. The number of training epochs (40) and the down-sampling strategy were identical to those used in Stage 2 fine-tuning. Distillation was performed using AdamW with a base learning rate of 1 × 10⁻³ and a weight decay of 0.05.

##### Training Datasets

Training datasets were curated from DIA and DDA datasets spanning diverse organisms, laboratories, and instruments. Dataset details, including sample species, instrument types, ProteomeXchange identifiers and references, are provided in Supplementary Table 2. None of the training datasets were used for benchmark evaluation.

### In silico spectral library predictor

DelPi constructs an in silico spectral library to provide theoretical peaks and predicted retention times for PmSM construction and scoring.

#### Fragment-Intensity Predictor

Fragment-ion intensities are predicted using a four-layer Transformer model. The model takes the peptide sequence, modification profile, precursor charge, and collision energy as inputs, and output predicted intensities for eight fragment-ion channels, corresponding to b and y ions at charge states 1 and 2 and their associated neutral-loss variants. Neutral losses of phosphoric acid (H₃PO₄) are considered when phosphorylation modifications are present in peptide sequences.

#### Retention-Time (RT) Predictor

Retention time is estimated using a hybrid sequence model that combines convolutional layers for local motif extraction with recurrent layers to model long-range dependencies along the peptide sequence.

#### Training data and Optimization

Both predictors were trained on the PROSPECT dataset^20^, a large collection of MS2 spectra and retention times for synthetic peptides. Approximately 2 million and 115 million training examples were used for the retention-time and fragment-intensity predictors, respectively. The RT predictor was trained for 40 epochs using the full training set, whereas the fragment-intensity predictor was trained for 50 epochs with 3 million examples randomly down-sampled per epoch. Training was performed using an MSE loss and the AdamW optimizer with a base learning rate of 1×10⁻³ and a weight decay of 0.05.

### Search Workflow

A two-stage search workflow is shown in Fig. 1c and Supplementary Fig. 1.

#### Spectral library generation

Protein sequences from a user-provided FASTA file are digested in silico (for example, trypsin). For target–decoy FDR estimation, decoy peptides are generated using either a pseudo-reverse strategy or a mutation-based method as implemented in DIA-NN. In the mutation-based approach, amino acids G/A/V/L/I/F/M/P/W/S/C/T/Y/H/K/R/Q/E/N/D are substituted with L/L/L/V/V/L/L/L/L/T/S/S/S/S/L/L/N/D/Q/E, respectively, and this method was used by default. For each peptide, the fragment-intensity and retention-time predictors generate predicted fragment intensities and retention times. Predicted retention times are calibrated at search time using polynomial regression between predicted and observed retention times, fitted using 1% FDR target matches within each dataset.

#### First search

##### DIA PmSM extraction

For DIA data, DelPi identifies chromatographic peak groups, defined as sets of fragment-ion signals that co-elute over a shared retention-time interval, for each candidate precursor. To limit the number of peak-group candidates, DelPi applies a simple heuristic scoring scheme based on fragment-ion co-elution and retains up to ten peak groups per precursor. The heuristic score is computed as the mean cubed Pearson correlation across fragment-ion extracted ion chromatograms (XICs), in which pairwise correlations are cubed to emphasize strongly co-eluting fragment pairs before averaging. For each peak group, DelPi collects nine consecutive MS2 spectra around the elution apex and pairs each MS2 spectrum with the nearest MS1 spectrum, yielding a PmSM comprising nine MS1 and nine MS2 spectra (Supplementary Fig 4. a).

##### DDA PmSM extraction

For DDA data, candidate MS2 spectra for a given precursor are selected using a wide isolation-window criterion: precursor m/z ∈ [*W*_*min*_ − 3.5/*z*, *W*_*max*_ + 1.25/*z*], where z is the precursor charge and *W*_*min*_ and *W*_*max*_ denote the lower and upper m/z bounds of the MS2 isolation window. Among these candidates, up to ten MS2 spectra with the highest numbers of matched fragment ions are selected. For each selected MS2 spectrum, DelPi constructs a PmSM by defining an elution window centered at the MS2 retention time and collecting spectra within this window. The width of the elution window is set to a representative chromatographic peak width estimated from MS1 data. To estimate this value, MS1 peaks are first grouped into bins of 1 m/z. For each bin, an intensity profile over retention time is constructed and chromatographic peaks are detected. The widths of these peaks are measured, and the median peak width across all bins is used as the representative chromatographic peak width. The elution window is then divided into nine equally spaced temporal bins; within each bin, an MS2 spectrum satisfying the isolation criterion is selected and paired with the nearest MS1 spectrum, yielding up to nine MS1 and nine MS2 spectra per PmSM (Supplementary Fig 4. b).

##### PmSM encoding and rescoring

Extracted PmSMs are encoded into latent embeddings and scored using the pre-trained PmSM encoder with an attached scoring head. Candidate PmSMs are filtered based on the raw logit output of the scoring head, and only candidates with logit values ≥ 1.0 are retained for subsequent semi-supervised rescoring.

For rescoring, DelPi trains a lightweight multilayer perceptron (MLP) classifier consisting of two fully connected layers with 64 and 32 neurons, respectively. The classifier is trained using PmSM embeddings together with an additional feature ΔRT, defined as the difference between the observed retention time and the calibrated predicted retention time. Following a Percolator-inspired procedure, decoy peptides (defined at the stripped peptide sequence level) are randomly partitioned into two equal subsets: one subset is used for MLP training together with target PmSMs, whereas the other subset is held out for FDR estimation.

The training uses a batch size of 512, four warm-up epochs, and up to 40 training epochs with early stopping (patience 3), together with a cosine learning-rate schedule and the AdamW optimizer (base learning rate 1 × 10⁻³, weight decay 0.01). The training data are split into 80% training and 20% validation sets, and early stopping is determined based on the validation loss. The same asymmetric focal loss used during fine-tuning is applied for rescoring. Because only half of the decoy PmSMs are used for training, decoy counts are multiplied by two when computing FDR.

##### Peak sharing and redundancy removal

PmSMs are clustered using a greedy procedure based on shared fragment peaks. Similarity is quantified using the Jaccard index, defined as the fraction of shared fragment peaks relative to the union of fragment peaks used for scoring. Each PmSM is assigned to the first existing cluster whose representative exceeds a similarity threshold of 0.6; otherwise, a new cluster is created. Only the highest-scoring PmSM per cluster is retained for downstream analysis.

#### Spectral library refinement via experiment-level transfer learning

Fragment-intensity and retention-time predictors are fine-tuned using high-confidence identifications from the first search (1% FDR), following an experiment-level transfer learning strategy similar to that used in AlphaDIA. Fine-tuning is performed for up to 40 epochs using the AdamW optimizer (base learning rate 1 × 10⁻⁴, weight decay 0.05) with early stopping (patience 3). The training data are split into 80% training and 20% validation sets, and early stopping is determined based on the spectral angle computed on the validation set. The refined spectral library is generated only for peptides whose PmSMs pass the minimum logit threshold in the first search.

#### Second search with refined library

DelPi repeats PmSM extraction and scoring using the refined spectral library. A logit cutoff of −1.0 is applied before semi-supervised rescoring.

#### Cross-run analysis

PmSM embeddings from all runs are pooled to train a global scoring model. The global scoring model uses the same architecture and training procedure as the rescoring model applied in the first search stage.

After global scoring, cross-run retention-time (RT) alignment is performed. The first run is treated as the reference run, and each remaining run is aligned independently to this reference. For each run pair, shared target PmSMs identified at 1% FDR are used to fit a linear regression model, which is then applied to compute aligned RT values for the non-reference run.

After RT alignment, DelPi estimates a representative RT for each precursor using kernel density estimation (KDE) on the distribution of aligned RTs across runs. Within each individual run, the PmSM whose RT is closest to this representative RT is selected as the final candidate for that precursor, resolving ambiguity when multiple candidate peak groups are present.

The selected PmSMs across all runs are used for global FDR estimation, which is performed using pooled scores and the split-decoy strategy described above. Precursor- and peptide-level FDRs are calculated. DelPi provides an internal protein inference module based on a greedy set-cover algorithm with heuristic grouping^21^, similar to the approach implemented in AlphaDIA. Protein inference can also be performed using external tools based on the reported PmSM scores. For benchmarking analyses, protein inference were performed using an external tool, PyProteinInference^22^ to ensure fair and consistent comparison across methods. Global protein-group-level FDRs are computed after protein inference.

##### Quantification for DIA data

Label-free quantification is performed at the fragment-ion level using a set of fragment ions consistent across runs. For each precursor, fragment ions are ranked by co-elution consistency with other fragments assigned to the same precursor. Within each run, co-elution consistency is computed as the average cubed Pearson correlation between the XIC of a fragment ion and the XICs of the other fragments. These scores are summed across runs, and the six fragments with the highest summed consistency are selected for quantification. Precursor intensities are computed by summing integrated peak areas of selected fragments. Peak areas are computed by trapezoidal integration over a fixed window of seven spectra (apex ±3 spectra). Protein-level quantification is performed using the MaxLFQ algorithm^23^.

### Benchmarking and evaluation

#### Benchmark datasets

Benchmarking was performed on four DIA datasets and two DDA datasets (Supplementary Table 1). The DIA datasets comprised (i) a mouse–yeast lysate dilution-series consisting of mixtures with defined mouse fractions (5–40%), used for both identification and DIA quantification benchmarking, (ii) a single-cell human proteome dataset, (iii) a neat human plasma dataset, and (iv) a mouse phosphoproteomics dataset. The DDA datasets were derived from CPTAC cohorts and consisted of (i) global proteomics data from colon adenocarcinoma (COAD) samples and (ii) TMT-labeled phosphoproteomics data from lung adenocarcinoma (LUAD) samples. For the neat plasma dataset and each DDA dataset, ten runs were randomly selected; the corresponding raw file names are provided in Supplementary Table 3. None of the benchmark datasets were used for model training, including both pre-training and fine-tuning.

#### Software and versions

For DIA benchmarks, DelPi was compared with DIA-NN ^3^ (v1.8.1), AlphaDIA^6^ (v1.12.1), and DIA-BERT^9^ (v1.1). For DDA benchmarks, comparisons were made with MS-GF+^4^ (v2024.03.26) and Sage^5^ (v0.14.7).

#### Search configuration

Search parameters were standardized across tools. The precursor m/z range was set to 300–1,800 m/z and the fragment m/z range to 200–1,800 m/z. Peptide length was restricted to 7–30 amino acids. Allowed precursor charge states were 1–4 and fragment charge states were 1–2. A maximum of one missed cleavage was allowed. Carbamidomethylation of cysteine was set as a fixed modification. For non-phosphoproteomics datasets, oxidation (M) and protein N-terminal acetylation were allowed as variable modifications (maximum 2). For phosphoproteomics datasets, phosphorylation (S, T, Y) was allowed as a variable modification (maximum 3). TMT labeling was treated as a fixed modification for the respective DDA datasets. All tools were configured to use 32 threads, and the FDR threshold was set to 1%.

For entrapment-based false discovery proportion (FDP) estimation, composite FASTA databases were constructed by merging the target-species proteome with an entrapment-species proteome (Supplementary Table 1).

Tool-specific configurations were as follows:

- **Sage:** wide-window search (precursor tolerance −3.5 to +1.25 Da) and chimeric search were enabled; up to five peptides reported per spectrum.
- **DIA-NN:** library-free mode with match-between-runs (MBR) enabled.
- **DIA-BERT:** spectral library generated by DIA-NN in library-free mode; peptides shared across multiple proteins were excluded.
- **AlphaDIA:** transfer-learning and transfer-library options enabled.

#### Hardware and computational constraints

Benchmarking was performed on a workstation equipped with two Intel Xeon Gold 6226R processors (32 cores total), 256 GB RAM, and an NVIDIA GeForce RTX 4090 GPU (24 GB VRAM). Due to memory and computational constraints, DIA-BERT was evaluated only on a single run of the mixed-species dilution-series dataset (HF_20210927_M200-Y800_DIA_2.raw), whereas AlphaDIA was excluded from both the full dilution-series analysis and the phosphoproteomics comparison.

#### Evaluation

Entrapment-based FDP was computed as described previously^19^, reporting both the lower-bound (LB) FDP and the combined (Comb) FDP. The LB FDP represents a conservative estimate of false discoveries, defined as the fraction of entrapment-species identifications among reported targets, and therefore provides a strict lower bound on the true FDP. The combined FDP further adjusts this estimate by accounting for the relative sizes of the target and entrapment databases, yielding a more realistic estimate of the overall false discovery proportion.

Global FDR handling differed across tools based on benchmarking scope and output characteristics. DIA-BERT was benchmarked on a single run and was therefore excluded from global FDR estimation; its native FDR estimates were used. For DIA-NN, native global FDR estimates were used, with ‘LIB.Q.Value’ and ‘Lib.PG.Q.Value’ serving as precursor- and protein-group–level global FDRs, respectively. For all other tools, precursor- and peptide-level global FDRs were derived from the reported confidence scores. For AlphaDIA, the posterior error probability reported as ‘precursor.proba’ was used, following the documented output specification. For MS-GF+, the ‘SpecEValue’ score was used, and for Sage, the ‘sage_discriminant_score’ was used to derive global FDRs.

Protein inference was standardized across tools using PyProteinInference^22^ when decoy match information was available. For DIA-NN and DIA-BERT, protein inference used native outputs because decoy match results required for external inference were not available. Performance was evaluated by comparing the number of identified precursors and protein groups at a 1% global FDR threshold.

#### Label-Free Quantification (LFQ)

LFQ performance was assessed using the dilution-series dataset. Analysis was restricted to protein groups composed exclusively of mouse proteins. A protein group was considered quantifiable if it was quantified in at least three of five replicates for a given mixing ratio.

## Data availability

All datasets used for model training and benchmarking are publicly available from PRIDE, MassIVE, or other community repositories. Accession numbers and download links are provided in Supplementary Tables 1 and 2.

## Code availability

The source code is also accessible at https://github.com/bertis-informatics/delpi under the MIT license. All benchmarking and analysis scripts used in this study are publicly available at https://github.com/bertis-informatics/delpi-benchmark.

## Acknowledgements

This work was supported by a National Research Foundation of Korea (NRF) grant funded by the Korea government (MSIT) (RS-2024-00454407).

## Author contributions

J.P. and S.K. designed the study, and J.P. developed and implemented DelPi and performed all benchmarking experiments. S.K. provided scientific supervision and guidance. K.K. contributed institutional support. U.K. provided administrative oversight and reviewed the manuscript. J.P. and S.K. wrote the manuscript with input from all authors.

