## Supplementary for "DelPi Learns Generalizable Peptide–Signal Correspondence for Mass Spectrometry-Based Proteomics"

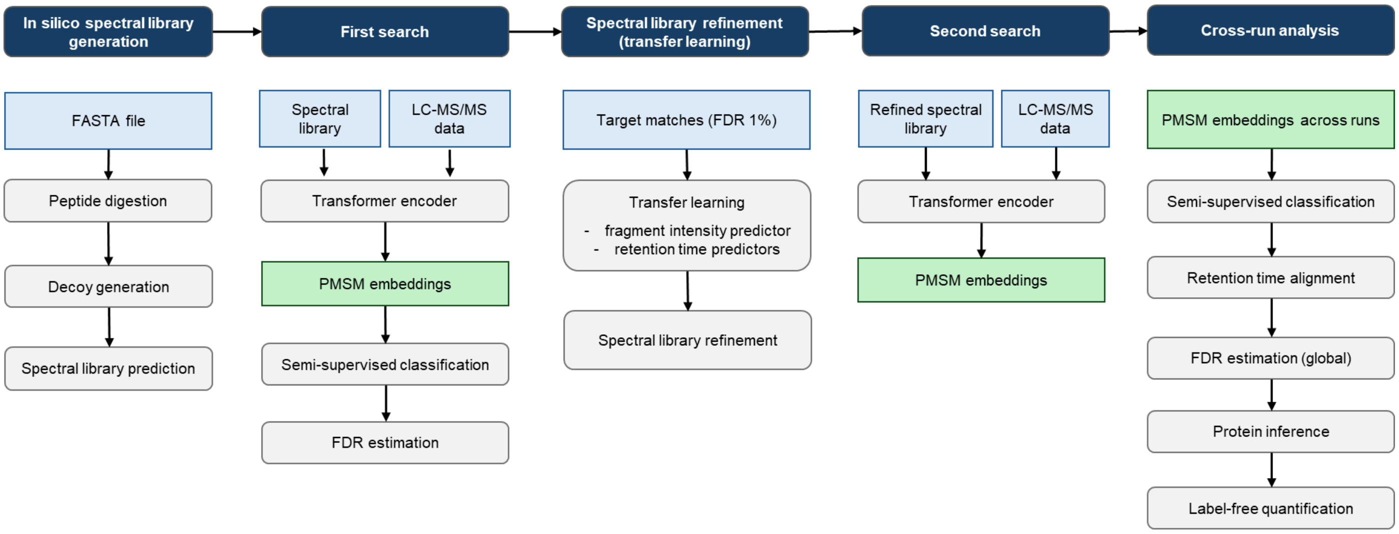


**Supplementary Figure 1. Detailed DelPi search workflow.** The DelPi search workflow follows a two-stage identification strategy with experiment-level adaptation and cross-run consolidation. Protein sequence databases are first processed by in silico digestion and decoy generation, followed by deep learning–based prediction of fragment-ion intensities and peptide retention times to construct an initial in silico spectral library. In the first search stage, given candidate peptides, Peptide–Multi-Spectra Matches (PmSMs) are extracted from liquid chromatography–tandem mass spectrometry (LC–MS/MS) data and encoded by the pre-trained Transformer encoder to generate PmSM embeddings, which are used for semi-supervised classification to identify high-confidence targets at 1% false discovery rate (FDR). These high-confidence identifications are then used for experiment-level transfer learning to adapt fragment-intensity and retention-time prediction models to the specific instrument and chromatographic conditions, yielding a refined spectral library. A second, more sensitive search is performed using the refined library to generate updated PmSM representations and identification scores. Finally, results are consolidated across runs through global scoring, retention-time alignment, protein inference, and label-free quantification.

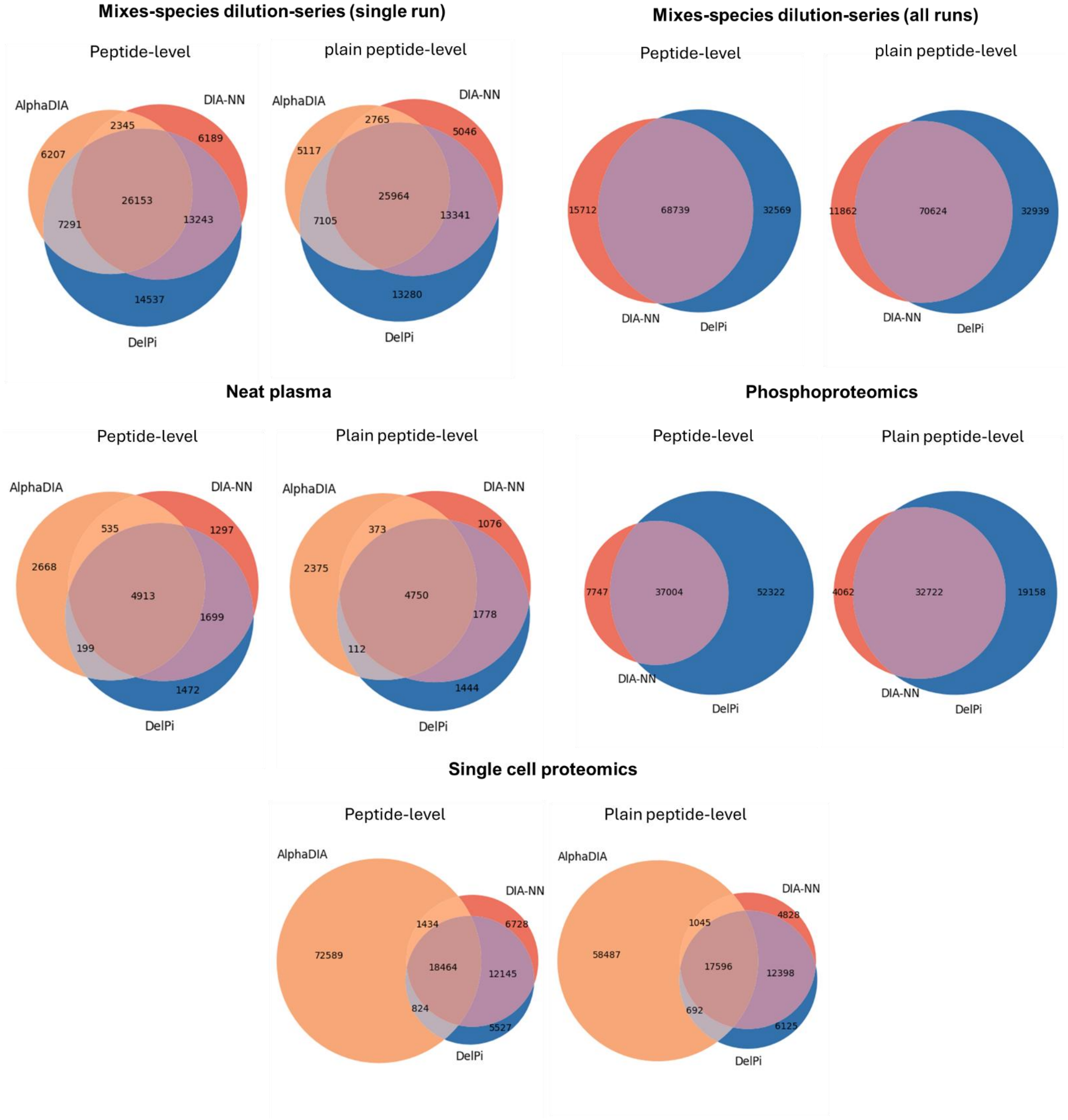


**Supplementary Figure 2. Overlap of peptides identified at 1% FDR in DIA datasets.** For each method, peptide identifications were filtered at 1% FDR. Because DIA-NN does not explicitly report peptide-level FDR, its identifications were filtered using a 1% precursor-level FDR for comparison. Overlaps were additionally evaluated using stripped peptide sequences (“plain peptides”), in which post-translational modification (PTM) annotations were removed. For the dilution-series dataset (single-run analysis), the Venn diagram includes only the three methods with the largest numbers of peptide identifications (DIA-NN, DelPi, and AlphaDIA); DIA-BERT was excluded from the Venn representation for clarity.

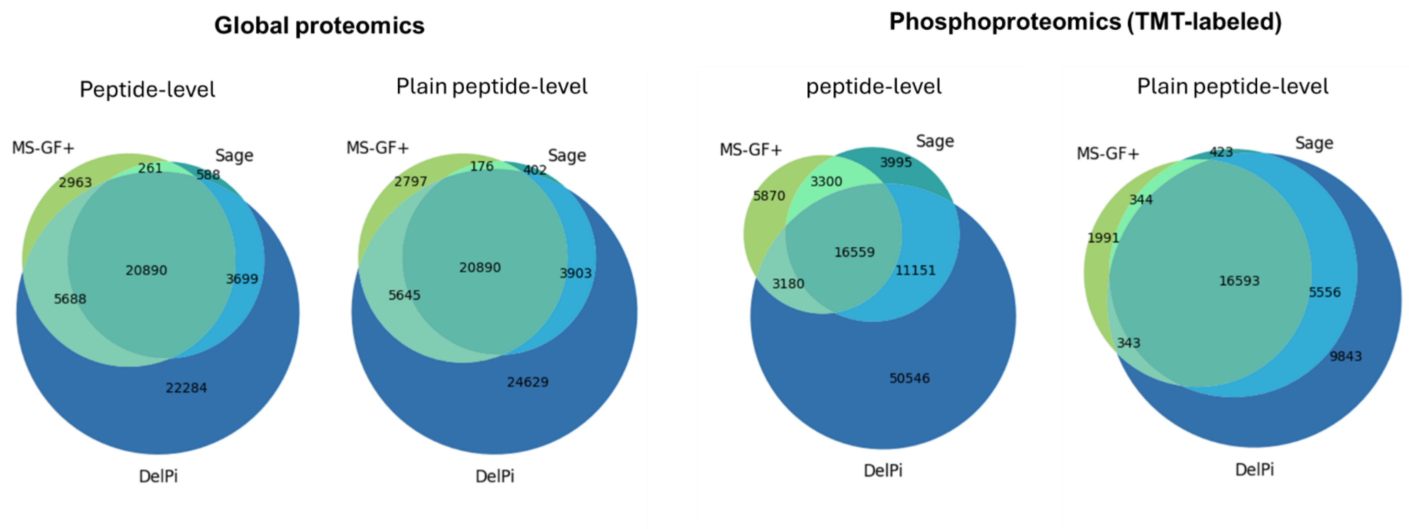


**Supplementary Figure 3. Overlap of peptides identified at 1% FDR in DDA datasets.** For each method, peptide identifications were filtered at 1% FDR. Overlaps were also evaluated using stripped peptide sequences (“plain peptides”), excluding PTM annotations.

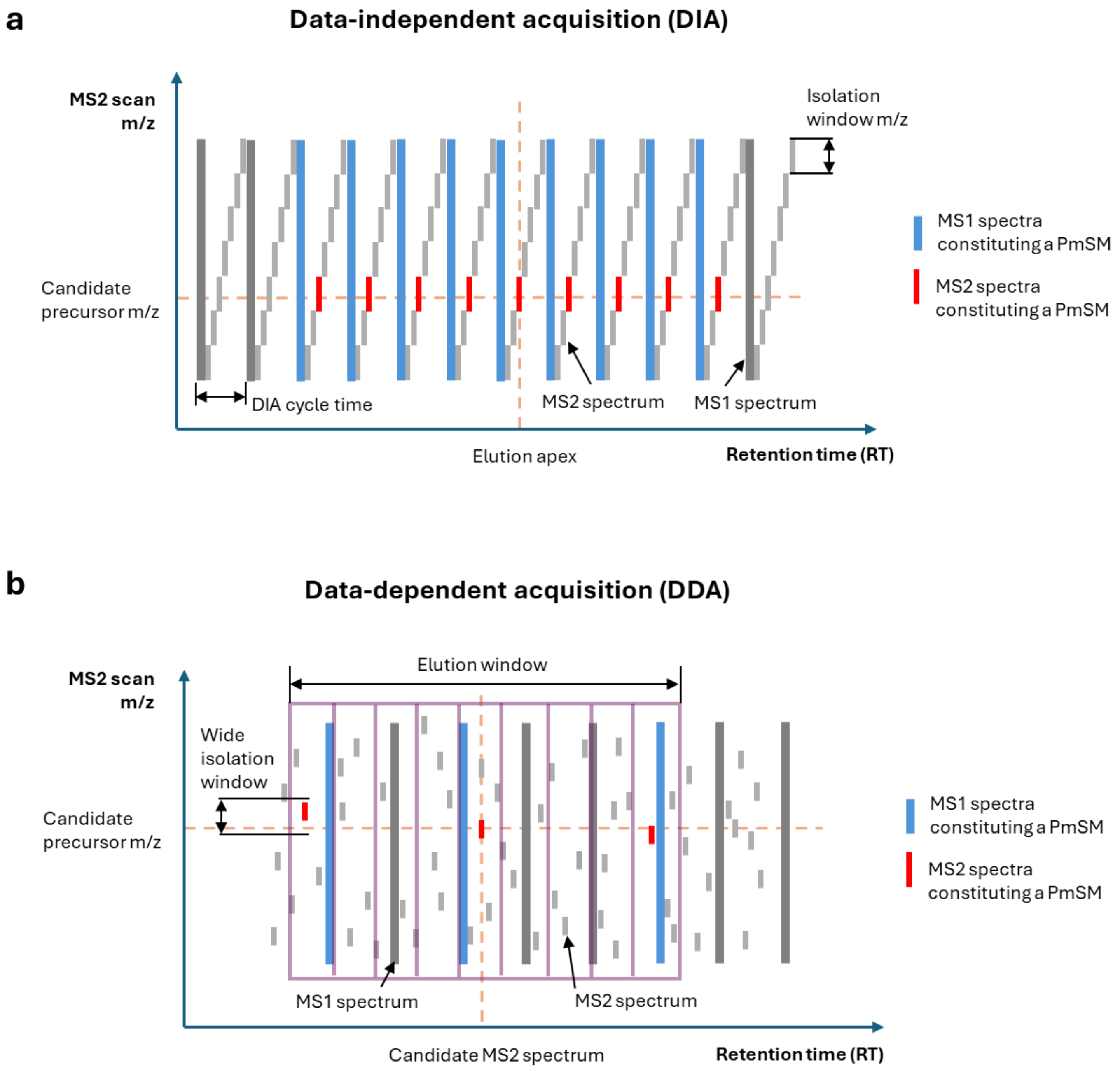


**Supplementary Figure 4. PmSM extraction for DIA and DDA data. a, DIA PmSM extraction.** Given a candidate chromatographic peak group for a precursor in DIA data, nine constituent MS2 spectra are collected symmetrically around the elution apex of the peak group. Each MS2 spectrum is paired with its nearest MS1 spectrum, yielding a Peptide–Multi-Spectra Match (PmSM) composed of nine MS1 and nine MS2 spectra. **b, DDA PmSM extraction.** Given a candidate MS2 spectrum for a precursor in DDA data, an elution window is defined by centering the window at the MS2 retention time, with the window width set to a representative chromatographic peak width estimated from MS1 data. The elution window is divided into nine equally spaced temporal bins. Within each bin, an MS2 spectrum satisfying a wide isolation-window criterion— precursor m/z $\in [W_{min} - 3.5/z, W_{max} + 1.25/z]$, where z is the precursor charge and $W_{min}$ and $W_{max}$ denote the lower and upper m/z bounds of the MS2 isolation window—is selected and paired with its nearest MS1 spectrum, yielding up to nine MS1 and nine MS2 spectra per PmSM.

**Supplementary Table 1. Benchmark dataset.**

| Name | Acquisition | Species | Entrapment | FASTA download | Instrument | Repository ID | Reference |
|---|---|---|---|---|---|---|---|
| Dilution series | DIA | Mouse, Yeast | Arabidopsis | 2022-03-02 | Q Exactive HF | PXD034709 | Lou et al., Nat. Commun., 2023 |
| Single cell | DIA | Human | E.coli | 2022-02-18 | Astral | PXD049412 | Bubis et al., Nat. Methods, 2025 |
| Neat Plasma | DIA | Human | E.coli | 2022-02-18 | Q Exactive HF-X | PXD038394 | Scott et al., Nat. Commun., 2023 |
| Phosphoproteomics | DIA | Mouse | Yeast | 2021-11-03 | Astral | MSV000093613 | Lancaster et al., Nat. Commun., 2024 |
| Global proteomics | DDA | Human | E.coli | 2022-02-18 | Q-Exactive Plus | PDC000109 | Vasaikar et al., Cell, 2019 |
| Phosphoproteomics (TMT) | DDA | Human | E.coli | 2022-02-18 | Fusion Lumos | PDC000149 | Gillette et al., Cell, 2020 |

**Supplementary Table 2. Training dataset.**

| Dataset | Species | Acquisition | Instrument | Repository ID | Reference |
|---|---|---|---|---|---|
| 2018-HeLa | Human | DIA | Q Exactive HF | MSV000082805 | Searle et al., Nat Commun, 2018 |
| 2020-Yeast | Yeast | DIA | Fusion Lumos Tribrid | MSV000084000 | Searle et al., Nat Commun, 2020 |
| lymph_ecoli | Human, E.coli | DIA | Eclipse | EGAD00010002223 | Fröhlich et al., Nat Commun, 2022 |
| ccRCC | Human | DIA | Fusion Lumos Tribrid | PDC000200 | Clark et al., Cell, 2019 |
| melanoma-phospho | Human | DIA | Fusion Lumos Tribrid | PXD022992 | Gao et al., Mol. Omics, 2021 |
| low-input-cell | Human | DIA | Eclipse | PXD027679 | Siyal et al., Anal. Chem., 2021 |
| single-cell | Human | DIA | Eclipse | PXD023325 | Gebreyesus et al., Nat. Commun., 2022 |
| Astral-HeLa | Human | DIA | Astral | PXD042704 | Heil et al., J. Proteome Res., 2023 |
| 2023-HeLa | Human | DIA | Astral | MSV000095138 | Wallmann et al., Nat. Biotechnol., 2025 |
| 2023-narrow-win | Human | DIA | Astral | PXD046357 | Guzman et al., Nat. Biotechnol., 2024 |
| | Yeast | DIA | Astral | PXD046386 | |
| | Human | DIA/DDA | Astral | PXD046453 | |
| 2021-Yeast | Yeast | DDA | Q Exactive Plus | PXD009815 | Bouyssié et al., Bioinformatics, 2020 |
| 2022-Plasma | Human | DDA | Exploris 480 | PXD037340 | Kverneland, et al., Proteomics, 2022 |
| 2023-Mouse | Mouse | DDA | Q Exactive HF | PXD035255 | Watanabe et al., Sci. Rep., 2023 |
| 2022-Jurkat | Human | DDA | Exploris 480 | PXD036024 | Zambo et al., Sci. Adv., 2023 |
| 2019-HeLa | Human | DDA | Exploris 480 / Q Exactive HF | PXD042233 | Webel et al., Sci. Data, 2024 |
| 2018-NCI7 | Human | DDA | Fusion Lumos / Q Exactive | PXD008952 | Clark et al., J. Proteome Res., 2018 |

**Supplementary Table 3. Randomly selected run files for benchmark test**

| Dataset | Randomly selected run files | Dataset | Randomly selected run files |
|---|---|---|---|
| Neat plasma (DIA) | TM_M2012_030.raw | Phosphoproteomics (TMT, DDA) | 24CPTAC_COprospective_W_VU_20150215_01CO001_f03.raw |
| | TM_M2012_054.raw | | 24CPTAC_COprospective_W_VU_20150215_01CO001_f05.raw |
| | TM_M2012_061.raw | | 38CPTAC_COprospective_W_VU_20150904_11CO031_f01.raw |
| | TM_M2012_095.raw | | 40CPTAC_COprospective_W_VU_20150909_05CO039_f02.raw |
| | TM_M2012_100.raw | | 49CPTAC_COprospective_W_VU_20151222_11CO062_f02.raw |
| | TM_M2012_110.raw | | 62CPTAC_COprospective_W_VU_20160311_05CO041_f04.raw |
| | TM_M2012_130.raw | | 69CPTAC_COprospective_W_VU_20160407_22CO006_f06.raw |
| | TM_M2012_137.raw | | 85CPTAC_COprospective_W_VU_20160630_11CO058_f05.raw |
| | TM_M2012_172.raw | | 88CPTAC_COprospective_W_VU_20160705_11CO054_f04.raw |
| | TM_M2012_189.raw | | 92CPTAC_COprospective_W_VU_20160728_11CO061_f02.raw |
| Global proteomics (DDA) | 08CPTAC_LUAD_P_BI_20180504_BD_f07.raw | | |
| | 09CPTAC_LUAD_P_BI_20180509_BD_f06.raw | | |
| | 12CPTAC_LUAD_P_BI_20180807_BD_f06.raw | | |
| | 13CPTAC_LUAD_P_BI_20180526_BD_f02.raw | | |
| | 13CPTAC_LUAD_P_BI_20180526_BD_f09.raw | | |
| | 18CPTAC_LUAD_P_BI_20180629_BD_f05.raw | | |
| | 19CPTAC_LUAD_P_BI_20180701_BD_f08.raw | | |
| | 20CPTAC_LUAD_P_BI_20180705_BD_f03.raw | | |
| | 22CPTAC_LUAD_P_BI_20180726_BD_f07.raw | | |
| | 25CPTAC_LUAD_P_BI_20180803_BD_f03.raw | | |
